# Digital Microbe, a genome-informed data integration framework for team science on emerging model organisms

**DOI:** 10.1101/2024.01.16.575828

**Authors:** Iva Veseli, Michelle A. DeMers, Zachary S. Cooper, Matthew S. Schechter, Samuel Miller, Laura Weber, Christa B. Smith, Lidimarie T. Rodriguez, William F. Schroer, Matthew R. McIlvin, Paloma Z. Lopez, Makoto Saito, Sonya Dyhrman, A. Murat Eren, Mary Ann Moran, Rogier Braakman

## Abstract

The remarkable pace of genomic data generation is rapidly transforming our understanding of life at the micron scale. Yet this data stream also creates challenges for team science. A single microbe can have multiple versions of genome architecture, functional gene annotations, and gene identifiers; additionally, the lack of mechanisms for collating and preserving advances in this knowledge raises barriers to community coalescence around shared datasets. “Digital Microbes” are frameworks for interoperable and reproducible collaborative science through open source, community-curated data packages built on a (pan)genomic foundation. Housed within an integrative software environment, Digital Microbes ensure real-time alignment of research efforts for collaborative teams and facilitate novel scientific insights as new layers of data are added. Here we describe two Digital Microbes: 1) the heterotrophic marine bacterium *Ruegeria pomeroyi* DSS-3 with >100 transcriptomic datasets from lab and field studies, and 2) the pangenome of the cosmopolitan marine heterotroph *Alteromonas* containing 339 genomes. Examples demonstrate how an integrated framework collating public (pan)genome-informed data can generate novel and reproducible findings.

## Introduction

Expanded access to the genomic data of microbial organisms has been transforming the way we approach microbiology research. Genome sequences are subsequently enhanced with knowledge from experimental, modeling, and field studies (e.g., ^1–4)^ with the goal of yielding insights into microbial physiology, ecology, and biogeochemistry. Yet because different research teams independently consolidate and curate genome-related information via ad hoc solutions, these diverse data streams have created challenges for interoperable analyses, especially in collaborative work. More generally, the lack of a framework for establishing consensus versions of genome-linked reference data hinders community coalescence around shared datasets. To extend the impact of curated and collated microbial data beyond a single research group, requirements are: 1) an established reference dataset^5^, which provides existing and updated knowledge in a standardized format; and 2) open access to these data, which allows multiple groups to collaboratively analyze and update the same genome and genome-linked information. The power of establishing a strategy for the open exchange of consensus microbial data linked to reference genomes for emerging model organisms, whether they are laboratory cultures or those reconstructed from metagenomes, is increasing as team science takes on growing roles in environmental and life sciences research.

Contemporary software solutions for the analysis and exchange of microbial genomes and associated ‘omics survey data can be broadly characterized into three groups: 1) online portals that provide a centralized location for uploading or downloading genomes and/or ‘omics datasets; 2) online portals with embedded applications that allow the user to choose from pre-selected genomes and/or ‘omics datasets or, in some cases, upload their own data for analysis; and 3) downloadable tools that enable local analysis of genomes and/or ‘omics data (Table 1). While they provide important services for individual research groups, these solutions do not necessarily maximize the efficiency of collaborative team science efforts. Typically, datasets are provided either as raw data or as highly-polished summaries, and intermediate data products for coordination of downstream analyses are not maintained. Moreover, most existing solutions are centralized, in which case data curation and platform maintenance falls on a single entity vulnerable to loss of funding, while data format, updates, and accessibility are not fully under the control of researchers. An alternative solution that partially solves the data sharing needs of collaborative team science efforts is anvi’o^6^ (https://anvio.org), an open-source software platform that can integrate a variety of data streams into interoperable, standalone SQL databases that can serve as collaborative data products^6^; however, anvi’o data products are not version-controlled. Inspired by the state-of-the-art technical opportunities offered by anvi’o, here we propose a general framework for the distribution and collaborative analysis of ‘omics datasets that is conducive to team science efforts. The ‘Digital Microbe’ (DM) concept describes features of a data product (#1-3) and a data implementation framework (#4-5) that:

1. Stores a genome sequence with sequence-linked information (e.g., curated gene calls, user-defined functional annotations, etc).
2. Supports additional layers of genome-associated data (e.g., genomic regions of particular interest, mutant strain availability, protein structures, etc).
3. Supports additional layers of experimental or environmental survey data, including intermediate analysis results of value to the research team (e.g., transcriptomic or proteomic activity across different experimental conditions, environmental distribution patterns through metagenomic or metatranscriptomic read recruitment analyses, etc).
4. Enables version-controlled addition of new data layers or curation of existing ones iteratively by any researcher.
5. Stores and enables the export of information in a universal format that is accessible to other programs and centralized or decentralized analysis platforms.

**Table 1.** Limitations of existing solutions for the sharing of ‘omics information.

| Category | Examples | Limitations |
| --- | --- | --- |
| Centralized databases<br>(data upload/download) | NBCI RefSeq<sup>95</sup><br>DiatOmicBase<sup>96</sup><br>ECMDB<sup>97</sup><br>ProPortal<sup>55</sup> | Restricted data types<br>Often specific to one type of organism<br>Updates after the initial data deposit can be limited |
| Online portals<br>(data upload + online analysis tools) | JGI IMG/M<sup>43</sup><br>KBase<sup>98</sup><br>Galaxy<sup>99</sup><br>PATRIC<sup>100</sup><br>CyVerse<sup>101</sup><br>RAST<sup>102</sup><br>SILVAngs<sup>103</sup><br>BV-BRC<sup>104</sup><br>Phycocosm<sup>105</sup> | May not enable sharing of data and results between users<br>Analysis workflows are often black boxes without transparent or changeable parameters<br>Dependence on the platform's computational resources can hinder high-throughput analyses<br>May not facilitate automation of analysis workflows |
| Downloadable tools<br>(local analysis with database snapshots) | Prokka<sup>106</sup><br>KofamScan<sup>18</sup><br>BLAST+<sup>107</sup><br>GToTree<sup>108</sup><br>Pathway Tools<sup>109</sup><br>DRAM<sup>110</sup> | May lack version control and/or easy mechanisms for updating database snapshot<br>Can be difficult to share data/results between users with different local versions of the database snapshot<br>Tool functionality is typically constrained to or tailored towards one kind of analysis |

We developed the Digital Microbe concept and its implementation in the National Science Foundation (NSF) Science and Technology Center for Chemical Currencies of a Microbial Planet (C-CoMP; https://ccomp-stc.org) consisting of a research team geographically distributed across 12 institutions. Our construction of Digital Microbes enabled Center members to simultaneously access, analyze and update experimental and environmental datasets for the Center’s two model marine bacterial species, *Ruegeria pomeroyi* DSS-3 and *Alteromonas macleodii* MIT1002, including diverse data types ranging from ‘omics surveys and environmental parameters to metabolic models and metabolomes. Here we demonstrate the feasibility of the Digital Microbe concept as a solution addressing widespread needs in the microbiology community for reproducible, integrated data products and we describe Digital Microbe data packages for each of C-CoMP’s model bacteria. The first Digital Microbe compiles knowledge of transcriptional response by *Ruegeria pomeroyi* DSS-3 gathered from 8 independent studies carried out between 2014 and 2023 (DOI: 10.5281/zenodo.7304959); the second describes an *Alteromonas* pangenome created by merging data from 339 isolate and metagenome-assembled genomes (DOI: 10.5281/zenodo.7430118).

## Results and Discussion

### Digital Microbe: Concept and Implementation

At its core, a Digital Microbe is a curated and versioned public data package that is (1) ‘self-contained’ (i.e., it can explain itself and its contents) and (2) ‘extensible’ (i.e., others can extend a Digital Microbe data package with additional layers of information coming from new experiments). The package consists of multiple datasets organized and linked through reference to the genome of a single microbe or the pangenome of a group of microbes (Figure 1). Data collection consolidates information such as gene annotations, coverage and other read-mapping statistics, and sample metadata. These data types can be flexible in scope and the extensibility of Digital Microbes via the programmatic addition of new ‘omics data types make them future-proof.

**Figure 1.**
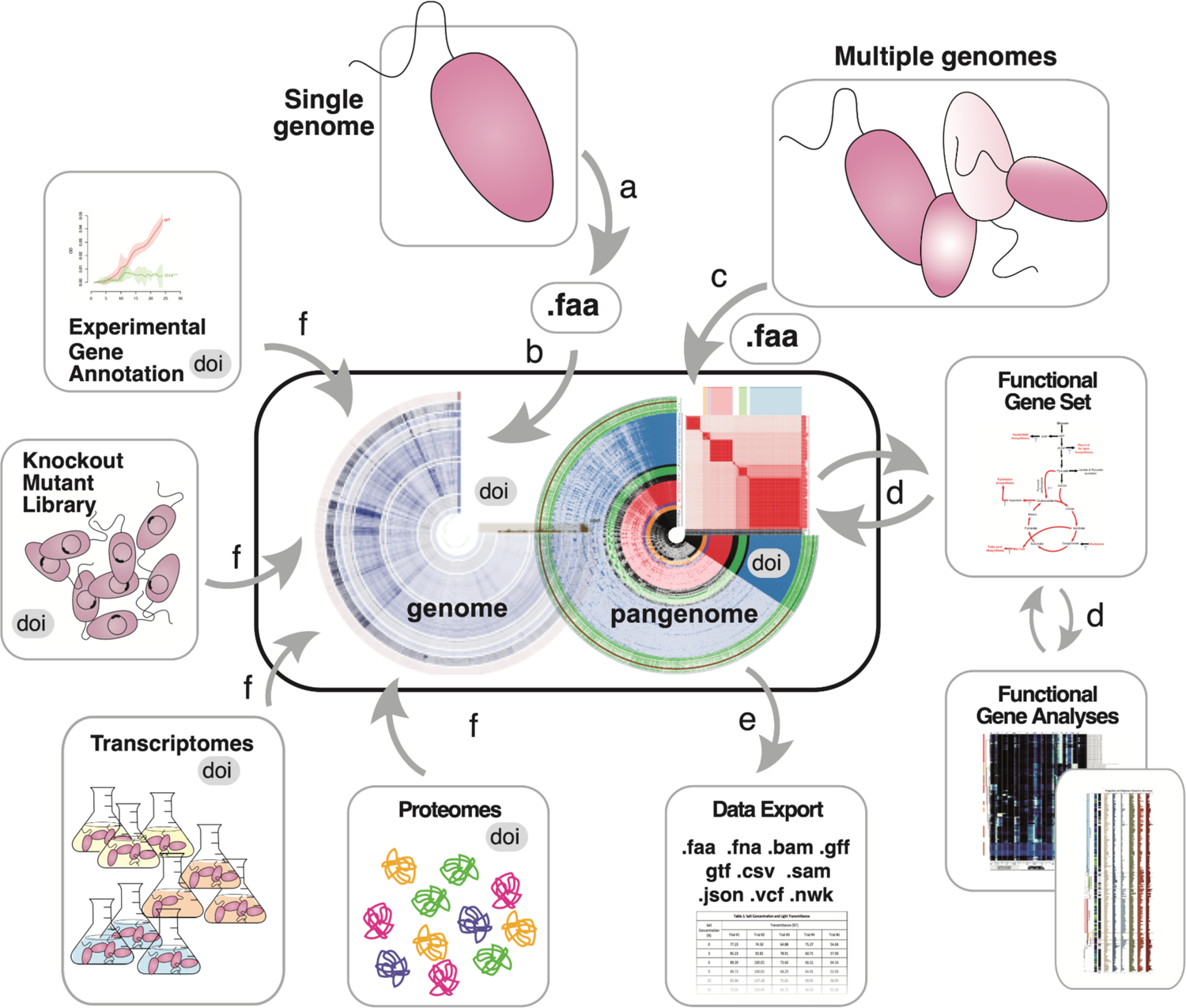
Architecture of a Digital Microbe. The genome of a model bacterium is (a) sequenced and (b) assembled and serves as the foundation of a Digital Microbe, a self-contained data package for a collaborative research team or a science community. (c) Alternatively, a pangenomic data package is assembled. (d) Intermediate datasets useful for downstream analyses are stored and reused, and (e) various data files and tables can be exported. (f) The Digital Microbe is iteratively populated with data layers referenced to individual genes, including mapped proteomes, transcriptomes, or gene-specific metadata types such as inventories of mutants or new annotations. Each Digital Microbe can be assigned a DOI (digital object identifier) and be versioned as new gene- or genome-referenced data are added.

The Digital Microbe framework utilizes a model organism’s genome or a clade’s pangenome as the foundation of a database file describing the DNA sequences (Figure 2). This database file is hosted in a central data repository where it can be accessed by collaborators and community members. A software platform was needed for collaborative analyses, and we chose the open-source software platform anvi’o^6^, which implements many of the Digital Microbe features described above (Figure 2). The concept behind the Digital Microbe framework, however, is independent of any one software platform. Similarly, C-CoMP hosts its Digital Microbe files on the data-sharing platform Zenodo (https://zenodo.org/), but other version-controlled storage solutions are available. As team science progresses, other genome- or gene-linked datasets (including both raw data and analysis results) can be added to the database by various groups, who update the publicly-hosted file to a new version that disseminates their data and findings to the team or community.

**Figure 2.**
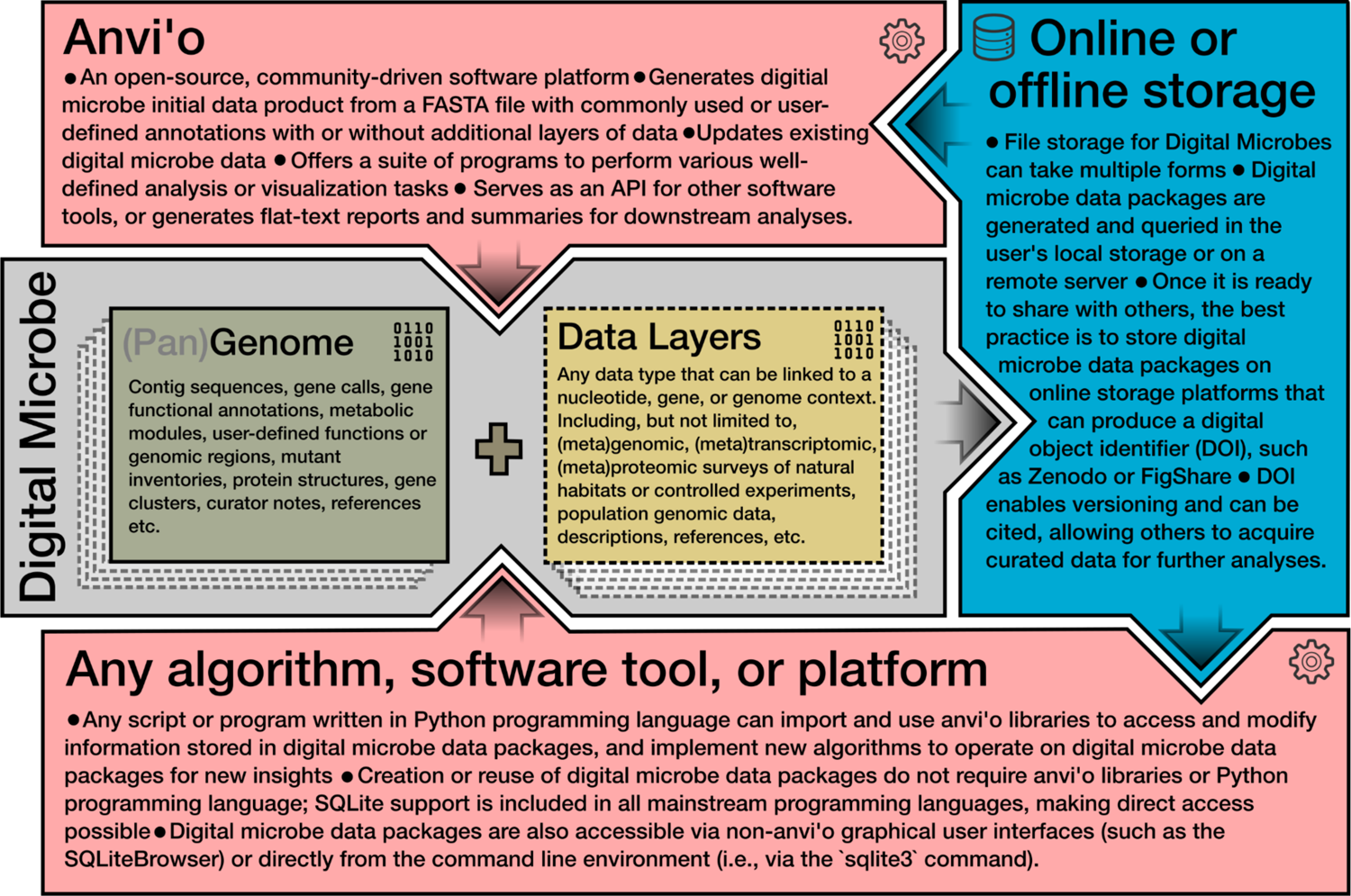
Situating the Digital Microbe concept in the existing computational environment. The Digital Microbe approach facilitates collaborative science by: establishing a version-controlled (pan)genomic reference; consolidating and cross-referencing collections of experimental and environmental data associated with a genome or pangenome; facilitating access to reusable intermediate analyses; and providing data export capabilities for transitioning to other programs or analysis software. While each of these features could be established by generating new software, we chose to use the existing open-source software platform anvi’o^6^, which implements several aspects of a Digital Microbe via (pan)genomic data storage in programmatically-queryable SQLite databases. The concept behind the Digital Microbe framework, however, is independent of any one software platform.

Here, we present two examples of Digital Microbes – one for the model organism *Ruegeria pomeroyi* and another for the pangenome of *Alteromonas* spp. – as well as case studies that exemplify how they can be used.

### The *Ruegeria pomeroyi* Digital Microbe

*Ruegeria pomeroyi* DSS-3 is a representative of the Roseobacteraceae family, an important bacterial group in marine microbial communities^7^ with its members among the most metabolically active bacterial cells in algal blooms and coastal environments^8^. *R. pomeroyi* has been well studied in the laboratory and field^9–11;^ it grows well in both defined and rich media; and it is amenable to genetic alteration^12,13^.

The *R. pomeroyi* Digital Microbe (Figure 3) is built on a well-curated genome assembly (*DM feature 1*) first annotated in 2004^14^, reannotated in 2014^15^, and enhanced with information from NCBI Clusters of Orthologous Groups (COG)^16^, Pfam^17^, and KEGG Kofam^18^. The Digital Microbe annotation is also continually updated (*DM feature 4*) with new experimental verifications of *R. pomeroyi* genes (e.g., ^15, 19–22)^ that have not been captured in standardized genome annotation repositories (e.g., RefSeq GCF_000011965.2). The *R. pomeroyi* DSS-3 Digital Microbe is available on Zenodo^23^.

**Figure 3.**
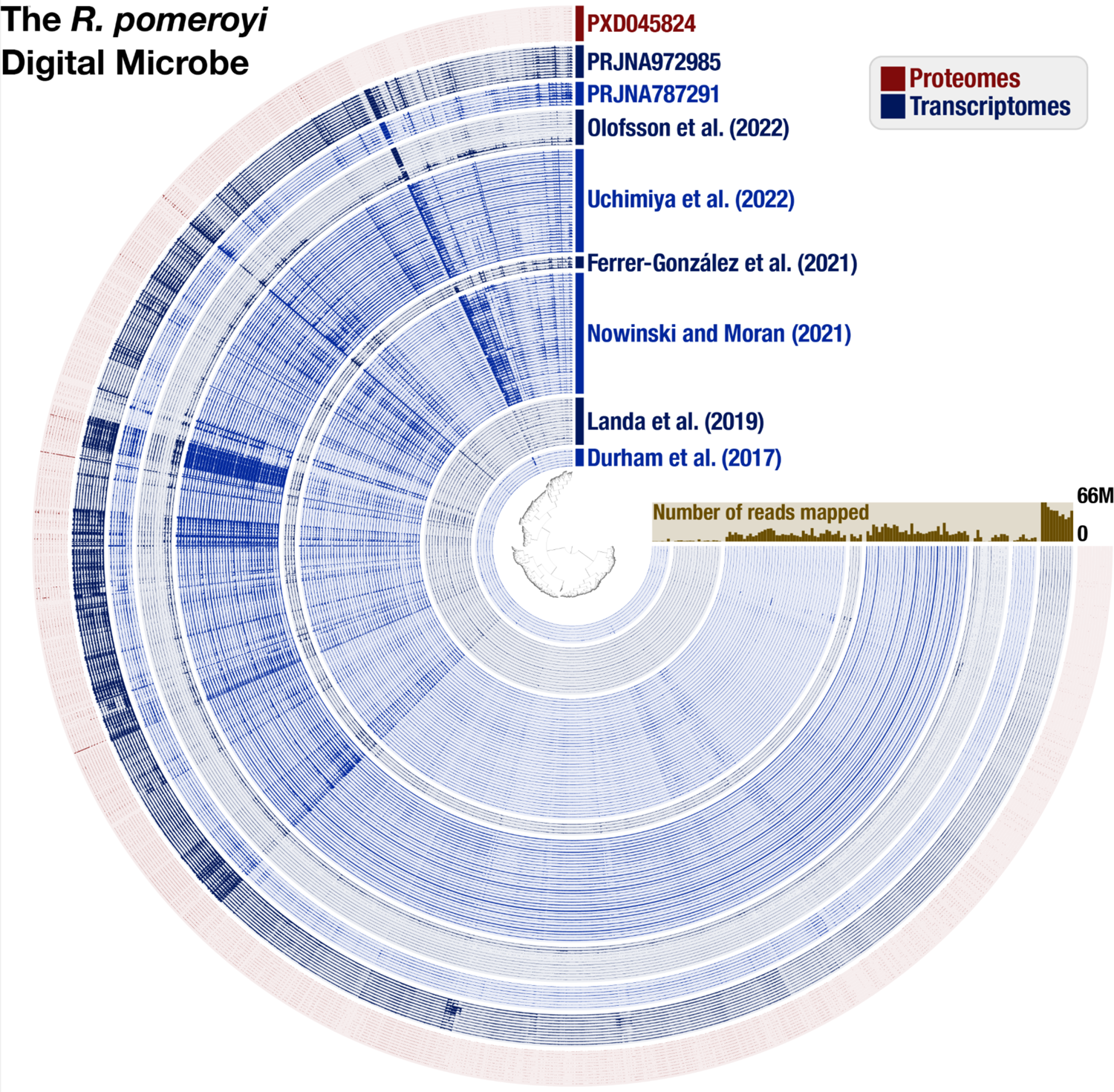
Contents of the *R. pomeroyi* Digital Microbe. As visualized in anvi’o ‘gene mode’, each item on the inner tree corresponds to one gene call in the *R. pomeroyi* genome, and the blue concentric circles display the coverage of each gene in a given transcriptome sample. The outermost red concentric circles correspond to normalized protein abundances from proteome samples (raw files available in the Proteomics Identifications Database (PRIDE) via Project PXD045824). Samples are grouped by their study of origin, with the data source indicated in text of the same color as the samples. The brown bar plot indicates the total number of reads that mapped from each transcriptome to the *R. pomeroyi* genome. This figure was generated from version 5.0 of the *R. pomeroyi* Digital Microbe databases on Zenodo.

#### A Use Case: Exploring the Substrate Landscape of R. pomeroyi

Our team is leveraging *R. pomeroyi* as a whole-cell biosensor of labile components of the marine dissolved organic carbon (DOC) pool. A recent study using *R. pomeroyi* knockout mutants definitively identified the cognate substrate of 18 organic compound transporters^24^ which were added to the Digital Microbe (*DM feature 2*). Previous homology-based annotations of most of these transporter systems were either incorrect or vague, and therefore of minimal ecological value. Although representing only a subset of the ∼126 organic carbon influx transporter systems in the *R. pomeroyi* genome, the presence or expression of these 18 is unequivocally linked to a known metabolite. With the new annotations in hand, we undertook a meta-analysis of transporter expression across 133 previously sequenced *R. pomeroyi* transcriptomes from laboratory and field studies between 2014 and 2023 to gain insights into the availability of these 18 metabolites in marine environments.

We added transcriptomes of *R. pomeroyi* to the Digital Microbe by mapping them onto the genome as individual data layers (Figure 3, Table S1) (*DM feature 3*). Using the anvi’o interactive interface, we established a custom dataset that consisted of the 62 protein components of the 18 experimentally annotated transporters (*DM feature 2*). We normalized the read counts for each protein to transcripts per million (TPM) and clustered the resulting data (Euclidean distance and Ward clustering). To generate a heatmap of transporter expression, we extracted the data from anvi’o and visualized it using python (*DM feature 5*).

This meta-analysis captured responses by *R. pomeroyi* to available substrates under 43 different ecological conditions (Figure 4), including during co-culture growth with phytoplankton^25–28^, on defined single or mixed substrates^20^, and after introduction into a natural phytoplankton bloom^10^. At the broadest scale, the transporters enabling organic acid uptake (acetate, citrate, fumarate, and 3-hydroxybutyrate) had the highest relative expression across conditions, together accounting for an average of 48% (range: 9.7 – 86%) of the transcripts for transporters with confirmed substrates. Recent studies have indeed discovered that Roseobacteraceae members are organic acid catabolic specialists^29,30^. Transporter transcription patterns also revealed the differences in substrate availability across environments. Introduced into a natural dinoflagellate bloom^10^, the citrate transporter had the highest relative expression; in a diatom co-culture, the acetate transporter was the most highly expressed; co-cultured with a green alga, transporter genes indicated that taurine, glycerol, carnitine, and dimethylsulfoniopropionate (DMSP) were on the menu. The organic acid transporter that enables *R. pomeroyi* uptake of 3-hydroxybutyrate^24^ was expressed across most growth conditions, yet this metabolite, also a precursor to the bacterium’s storage polymer polyhydroxybutyrate (PHB), has not previously been identified as a relevant currency in bacterially-mediated carbon flux. The meta-analysis also showed a pattern in expression for transporters that contain a substrate binding protein gene (i.e., the ABC and TRAP transporter classes): the gene is expressed at consistently higher levels than other genes in the same transporter (i.e., higher than permeases and ATP-binding proteins) despite all having membership in the same operon. Additional layers of gene regulation are therefore occurring either as within-operon differential expression or as post-transcriptional selective degradation. Regardless, this regulatory strategy would benefit a bacterium in an environment where substrate acquisition is the growth-limiting step.

**Figure 4.**
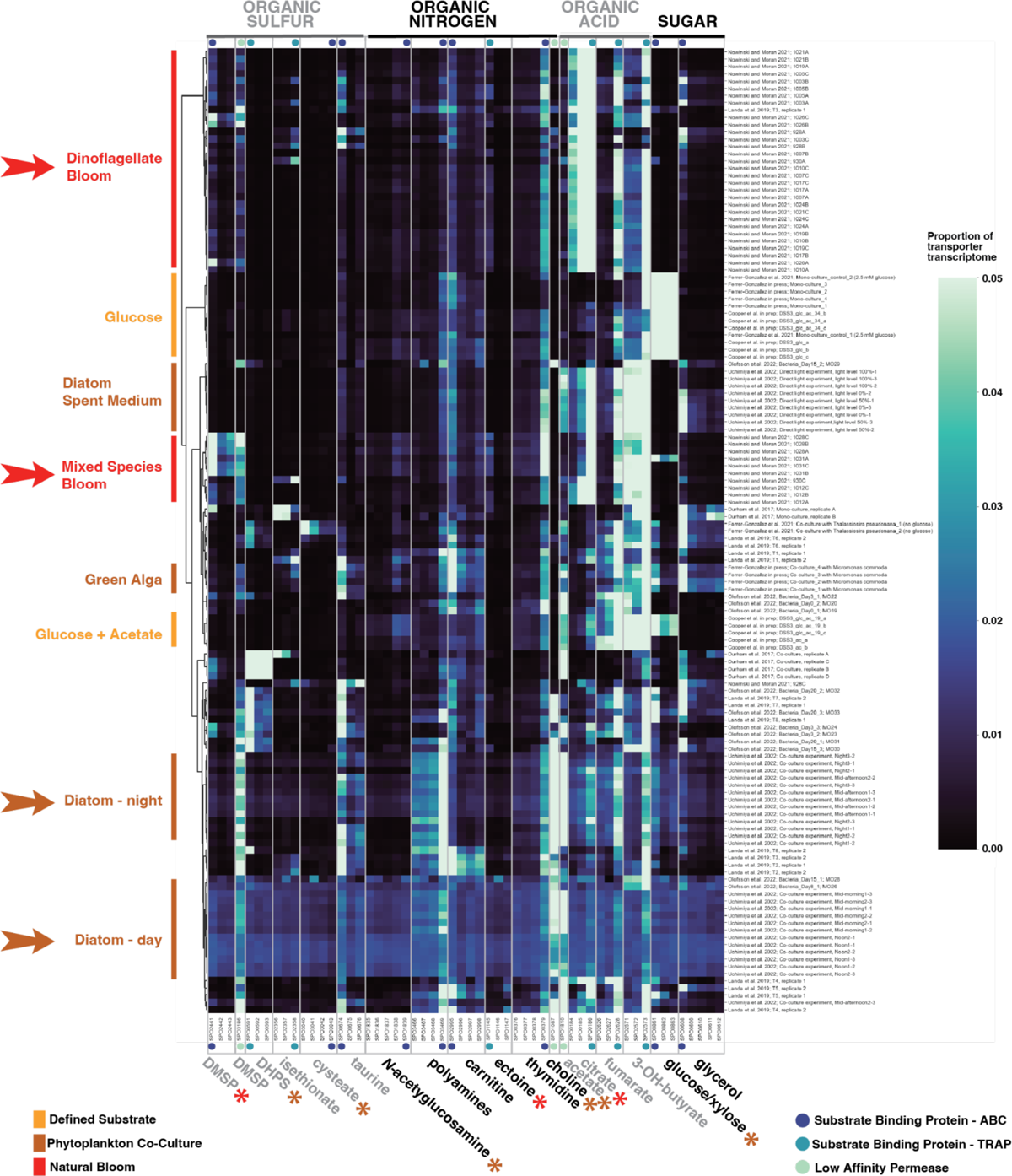
Clustered heatmap of relative gene expression for 18 experimentally annotated *R. pomeroyi* transporters compiled in a Digital Microbe. Each row represents a single transcriptome from the Digital Microbe dataset, and each column represents all transporter proteins with experimentally confirmed cognate substrates. Row labels indicate study and sample name (Table S1). Brighter colors indicate higher proportional expression (the scale maximum is ≥5% of the sum of the 18 transporter transcriptomes) while darker colors indicate lower. Arrows point out transcriptomes collected when substrates were derived from dinoflagellate-rich natural communities (red) or diatom co-cultures (brown); significant differences in transporter protein expression between these two substrate sources are indicated with asterisks colored red (enriched with dinoflagellates) or brown (with diatoms) (T-test, p≤0.05).

Three additional genome-linked data types recently added to the *R. pomeroyi* Digital Microbe include the locations of insertion sites of knockout mutants (covering 3,570 genes of 4,288 genes)^13,24^; proteomic data collected concurrently with one of the transcriptomic studies^31,32^, and TnSeq mutant fitness measurements in synthetic microbial communities^11^ (*DM feature 2*); these are enhancing collaborations among team members.

### The *Alteromonas* Digital Microbe

*Alteromonas* is a genus of marine Gammaproteobacteria whose members associate with particles and can contribute significantly to heterotrophic activity of phytoplankton blooms, sometimes in the role of helper bacteria that provide benefits to the phytoplankton^33–35^. Bacteria in this genus are distinguished by genomes encoding an average of 4,000 genes that enable use of a broad spectrum of substrates^36^, provide protection from reactive oxygen species to community members^33^, and mediate polysaccharide degradation^37^. The type species of the genus is *Alteromonas macleodii*^34,38^, with other notable species including *A. mediterranea*^39^, *A. australica*^40^, and *A. stellipolaris*^41^. While no single species has emerged as the primary model organism for this genus, the many genomes available for study provide an opportunity for pangenomic analysis to improve understanding of the evolution and diversity of this ubiquitous marine clade^42^.

The assembled pangenome consists of 336 *Alteromonas* genomes with genes called and annotated using one standardized pipeline (*DM feature 1*). Of these, 78 are isolate genomes^43–45^ and 258 are metagenome-assembled genomes (MAGs) obtained from a variety of marine environments in the global ocean^46^. Genomes represent members of the closely related ‘surface’ species *A. macleodii* (n=139) and ‘deep’ species *A. mediterranea* (n=25)^39^, and the widely distributed *A. australica* (n=63)^47^. The 34,390 gene clusters of the pangenome are linked to an imported phylogenetic tree assembled from single-copy core genes (see Methods), annotated using NCBI COGs^16^, KEGG KOfams^18^, CAZyme HMMs^48^ and orthology predictions from EggNOG-mapper^49–51^ (DM *feature 2*), and assigned as core or accessory genes for the genus (*DM feature 3*) based on a Bayesian approach available in anvi’o^52^. The *Alteromonas* Digital Microbe with relevant pangenome and phylogeny files is accessible on Zenodo^53^. Examples of future versioned enhancements of this Digital Microbe might include additions of new *Alteromonas* genomes and improved annotations from culture studies and novel annotation programs.

#### A Use Case: Evolutionary Patterns of Carbohydrate Use

We leveraged the information contained within the Digital Microbe to examine diversity in the ability of this opportunistic marine genus to use poly-/oligosaccharides^36^. Sugars and sugar polymers are an abundant and diverse component of the ocean’s dissolved organic carbon inventory^54^, and differences in how microbes use them provide important clues on the evolutionary diversification of their roles in the oceanic carbon cycle. Moreover, the ability to annotate genes with the Carbohydrate-Active enZYme (CAZyme) Database^48^ was recently added to anvi’o, allowing augmentation of the Digital Microbe with CAZyme annotations. The results indicate distinct CAZyme distributions across *Alteromonas* clades (Figure 6). For example, the *A. australica* and *A. stellipolaris* clades have more polysaccharide lyases than neighboring clades, while the *A. stellipolaris* clade is enriched in several other CAZyme categories as well. As patterns of diversity in CAZyme inventories are most distinct at the clade level compared to the within-clade level, carbohydrate utilization emerges as a potentially key driver of the large-scale niche partitioning of *Alteromonas* species.

**Figure 5.**
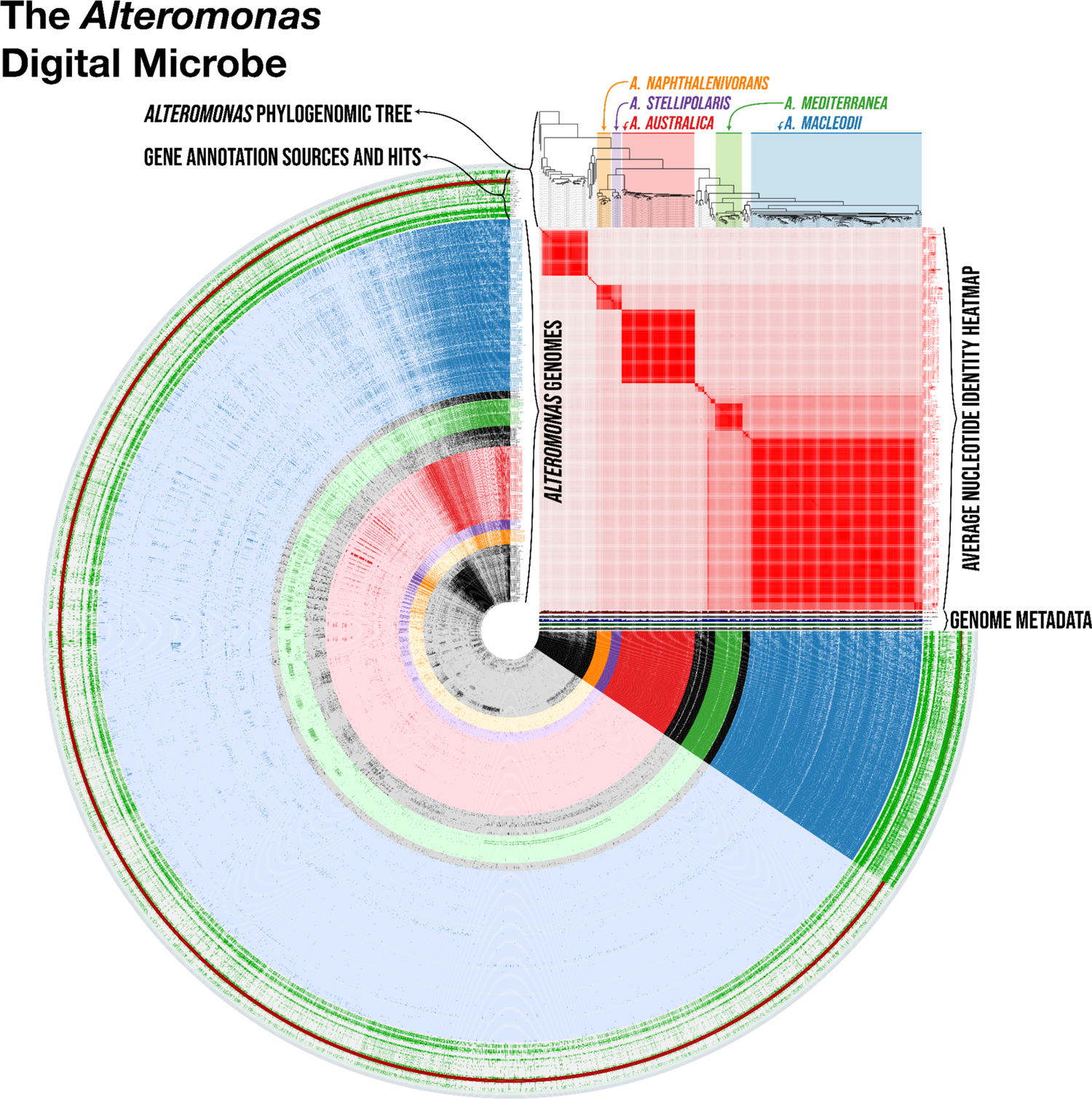
The *Alteromonas* Digital Microbe. Each concentric ring represents one *Alteromonas* genome, with colored rings identifying genomes from five clades of interest (*A. macleodii*, *A. mediterranea*, *A. austalica*, *A. stellioolaris*, and *A. naphthalenivorans*). The outermost green rings depict annotation sources applied to all genomes. Each spoke in the figure represents one gene cluster in the pangenome, with presence/absence denoted by darker/lighter colors, respectively. Genome metadata is shown next to each ring and includes total genome length, GC content, completion, number of genes per kbp, and number of gene clusters per genome. The red heatmap above the metadata shows the average nucleotide identity (ANI) percentage scores between genomes. The tree above the ANI heatmap shows the imported phylogenomic tree, with clades of interest color-referenced in the circular portion of the figure. This figure was generated using the anvi’o ‘anvi-display-pan’ from a version of the *Alteromonas* digital microbe without singleton genes, which is available on Zenodo under <u>DOI:10.5281/zenodo.10421034</u>.

**Figure 6.**
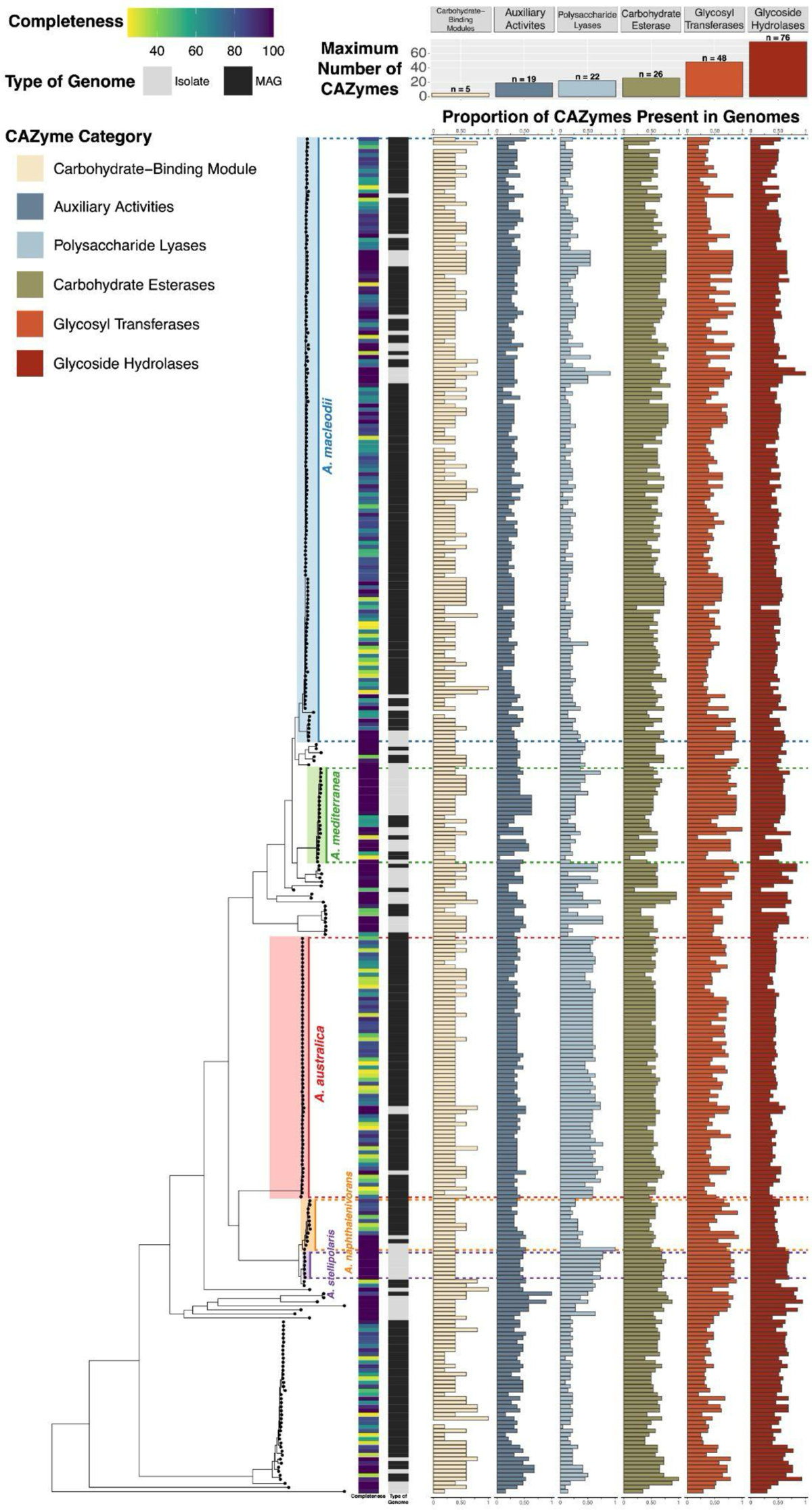
Distribution of CAZYme annotations across a phylogeny of 336 isolate and MAG genomes from the genus *Alteromonas*. The phylogeny of the genus is displayed on the left side of the figure, with genomes represented by points on the tree and five of the clades (*A. macleodii*, *A. mediterranea*, *A. australica*, *A. napthalenivorans*, and *A. stellipolaris*) highlighted. Each row on the right side of the figure represents one genome. Completeness and type of genome are shown in the two heatmaps to the right of the phylogeny. The horizontal bar plots of different colors show the proportion of CAZymes in each genome relative to the maximum number of all categories of CAZymes as indicated in the legend in the inset at the upper left. The maximum number for each CAZyme category is represented by the vertical bar plot at the top of the figure.

We also gained insights into how carbohydrate usage has shaped *Alteromonas* evolution and ecology from gene phylogenies of selected CAZymes (Supplementary Figure 1). The topology of several CAZyme phylogenies broadly recapitulates the topology of the genome phylogeny built from single-copy core genes (Supplementary Figure 2), suggesting that vertical descent has dominated the evolution of these genes. However, the topologies of other CAZyme phylogenies have significant discordance with the genome phylogeny (Supplementary Figure 2), suggesting that horizontal transfer has also had an important role in the evolution of carbohydrate utilization strategies in *Alteromonas*. The divergent evolutionary trajectories of different CAZymes highlight selective pressures acting on the metabolic diversification of *Alteromonas*, and may offer clues on how this diversification has in turn impacted the evolution of carbon cycling in the ocean.

### Future directions

Digital Microbe data packages furnish an architecture for reproducible, open, and extensible collaborative work in microbiology and its many derivative fields. While we present here a specific architecture tailored to our research focus, it is only one manifestation of the broader digital microbe concept: that decentralized taxon-specific databases are key mechanisms for capturing knowledge accumulating from genome-informed data that are now so vast and distributed as to be intractable to synthesize^55^. Digital Microbe packages allow one-stop shopping for data spread across multiple public archives, allow coordinated selection and documentation of genome structure and annotations within and between research teams, and are extensible to new data types. While the framework presented here is designed for bacterial and archaeal data, the development of digital microbes for eukaryotic model organisms is an important future application^56^. One enhancement under development by C-CoMP is an integrated toolkit for metabolic modeling, but the nature and scope of future applications can be defined by any research group that uses a digital microbe framework for their research. Organized and open access to taxon-explicit data is an essential foundation for modern microbiology.

## Methods

Both Digital Microbes were generated and analyzed using v7.1-dev or later versions of anvi’o^6^.

### Creation of the *Ruegeria pomeroyi* Digital Microbe

We created the *Ruegeria pomeroyi* Digital Microbe from the *R. pomeroyi* DSS-3 complete genome and megaplasmid sequences^14^ and (meta)transcriptome samples from ^10,27,28,32,57–59^. We generated a contigs database from the genome and megaplasmid sequences with ‘anvi-gen-contigs-databas’ and annotated the resulting Prodigal^60^ gene calls with *de novo* annotations from NCBI Clusters of Orthologous Genes (COGs)^16^, KEGG KOfams^18^, and Pfam^17^ via the associated anvi’o program for each annotation source. We also identified single-copy core genes using ‘anvi-run-hmms’ and associated these genes with taxonomy data from the GTDB^61^ using ‘anvi-run-scg-taxonomy’. We imported manually curated gene annotations, including annotations indicating which genes have available mutants^13^, using the program ‘anvi-import-functions’.

To process the (meta)transcriptomes, we quality-filtered the samples using FASTX-toolkit^62^ with the parameters described in ^25^. We mapped the reads to the DSS-3 genome using Bowtie 2^63^ and samtools^64^. Each sample’s read mapping data were converted into an anvi’o profile database using ‘anvi-profil’, and all samples were merged into a single database with ‘anvi-merg’. To add proteomic data^31^, we normalized spectral abundance counts with a normalized spectral abundance factor to make data comparable across all proteomes. We generated a ‘genes database’ to store gene-level information by running ‘anvi-interactiv’ on the established contigs and profile databases with the ‘--gene-mod’ flag, and we imported the normalized abundances for each gene into the genes database using the program ‘anvi-import-misc-datà. We also used this program to import fitness data associated with gene mutants from ^11^ into the same genes database.

### Transporter expression analysis for *Ruegeria pomeroyi*

The genes database in the *R. pomeroyi* Digital Microbe contains gene-level transcript coverage information from >100 samples. To assess the proportional expression of substrate-confirmed transporter genes, we used the anvi’o interactive interface to create a bin containing the transporter genes, and generated a static summary page with the “init-gene-coverages” box checked to export annotation and coverage data for each contig region where our genes of interest were located. After reading the exported data into dataframes using python v3.7.8 and pandas^65,66^, we extracted the coverage data for our specific genes of interest, normalized the coverages to TPM using the total number of reads per sample, and relativized these data to represent the proportional expression of each gene. We visualized these data as a clustermap using the seaborn package^67^ and assessed statistical differences in the mean gene expression using the a t-test implemented in the scipy stats package^68^.

### Creation of the *Alteromonas* Digital Microbe

To create the *Alteromonas* Digital Microbe, we collected 78 isolate genomes and 258 MAGs from the Joint Genome Institute’s Integrated Microbial Genomes (IMG) project^43^, NCBI^69^, and ^46^. We converted each genome into an anvi’o contigs database using ‘anvi-gen-contigs-databas’. For the genomes from IMG and NCBI, we determined completion and contamination statistics using CheckM v1.0.18^70^; for the MAGs that were taken from ^46^, we used the mean completeness and mean contamination statistics reported in that publication. We annotated the genes in each contigs database with the NCBI Clusters of Orthologous Genes (COGs)^16^, KEGG KOfams^18^, and Carbohydrate-Active enZYme (CAZyme) HMMs^48^ via the associated anvi’o program for each annotation source, and imported externally-run annotations from EggNOG-mapper^49–51^ and KEGG GhostKOALA^71^ into the databases using ‘anvi-import-functions’.

We ran ‘anvi-pan-genom’ to create the pangenome and computed the average nucleotide identity (ANI) between all pairs of genomes using ‘anvi-compute-genome-similarity’. To extract the core genome from the pangenome (i.e., genes found in all genomes), we used a Bayesian statistical method^52^ implemented in ‘anvi-script-compute-bayesian-pan-cor’. This method employs mOTUpan.py to determine the gene clusters likely to be core based on individual genome completeness scores.

### Phylogenomic Analysis of the *Alteromonas* genomes

To build the phylogeny of *Alteromonas*, we aligned and concatenated the sequences from 110 single-copy core gene clusters using ‘anvi-get-sequences-for-gene-clusters’. We imported these sequences into the tree building software RAxML, version 8.2.12^72^, and built the tree under the “PROTGAMMAAUTO” model setting. We used FigTree v1.4.4^73^ to midpoint root the tree and save it in newick file format. To incorporate the tree into the pangenome, we imported the newick tree with the program ‘anvi-import-misc-datà. For the phylogenomic CAZyme analysis, we used ‘anvi-split’ to subset gene clusters with known CAZyme functions into a new pangenome database and ran ‘anvi-summariz’ on this smaller pangenome to count the number of CAZymes per category, per genome. We visualized these data as a function of the previously-determined phylogeny in R v4.1.1^74^ using the packages aplot v0.1.9^75^, BiocManager v1.30.20^76^, dplyr v1.1.0^77^, ggnewscale v0.4.8^78^, ggplot2 v3.4.1^79^, ggstance v0.3.6.9000^80^, ggtree v3.7.1.003^81–85^, ggtreeExtra v1.9.1.992^81,86^, nationalparkcolors v0.1.0^87^, plyr v1.8.8^88^, RColorBrewer v1.1-3^89^, scales v1.2.1^90^, and tidyr v1.3.0^91^.

We then repeated the initial steps above to generate a phylogeny for the subset of isolate genomes (n=78), which resulted in a tree built from 111 single-copy core gene clusters. After subsetting the gene clusters with known CAZyme annotations into a smaller pangenome, we identified eight CAZyme-related gene clusters that were part of the single-copy core genome. We then generated an individual phylogeny from each of these CAZymes. We used R to compare the CAZyme phylogenies with the overall core genome phylogeny for these isolate genomes, with the packages listed above in addition to colorBlindness v0.1.9^92^, easyalluvial v0.3.1^93^, and gridExtra v2.3^94^.

## Data Availability

The *Ruegeria pomeroyi* Digital Microbe is available via DOI:10.5281/zenodo.7304959 and the *Alteromonas* Digital Microbe is available via DOI:10.5281/zenodo.7430118. The raw proteomics data included in the *Ruegeria pomeroyi* Digital Microbe is available on the Proteomics Identifications Database (PRIDE) project PXD045824 with accompanying metadata and processed data available in Biological and Chemical Oceanography Data Management Office (BCO-DMO) dataset 927507 via DOI: 10.26008/1912/bco-dmo.927507.1. The accompanying raw transcriptomic expression data to the proteomics data is available under the National Center for Biotechnology Information (NCBI) BioProject PRJNA972985 with metadata available in BCO-DMO dataset 916134 via DOI:10.26008/1912/bco-dmo.916134.1.

## Code Availability

Reproducible workflows for the generation of the Digital Microbes and the analyses described in this work can be accessed at https://github.com/C-CoMP-STC/digital-microbe. In particular, the Jupyter notebook for the *Ruegeria pomeroyi*use-case analysis can be found at https://github.com/C-CoMP-STC/digital-microbe/blob/main/rpom/rpom_dig_micro_transporter_expression_use_case.ipynb and the workflow for the *Alteromonas* use-case analysis can be found at the following link: https://github.com/C-CoMP-STC/digital-microbe/blob/main/alteromonas/useCase/alteromonasUseCases.md.

## Acknowledgements

This work was supported under NSF grant OCE-2019589 to the Center for Chemical Currencies of a Microbial Planet. IV acknowledges support by the National Science Foundation Graduate Research Fellowship under Grant No. 1746045. MAM acknowledges support by Simons Foundation Grant 542391 within the Principles of Microbial Ecosystems Collaborative. This is C-CoMP publication #024.

## Supplementary Tables

**Table S1.** Transcriptome samples included in the *Ruegeria pomeroyi* Digital Microbe.

## Supplementary Figures

**Supplementary Figure 1.**
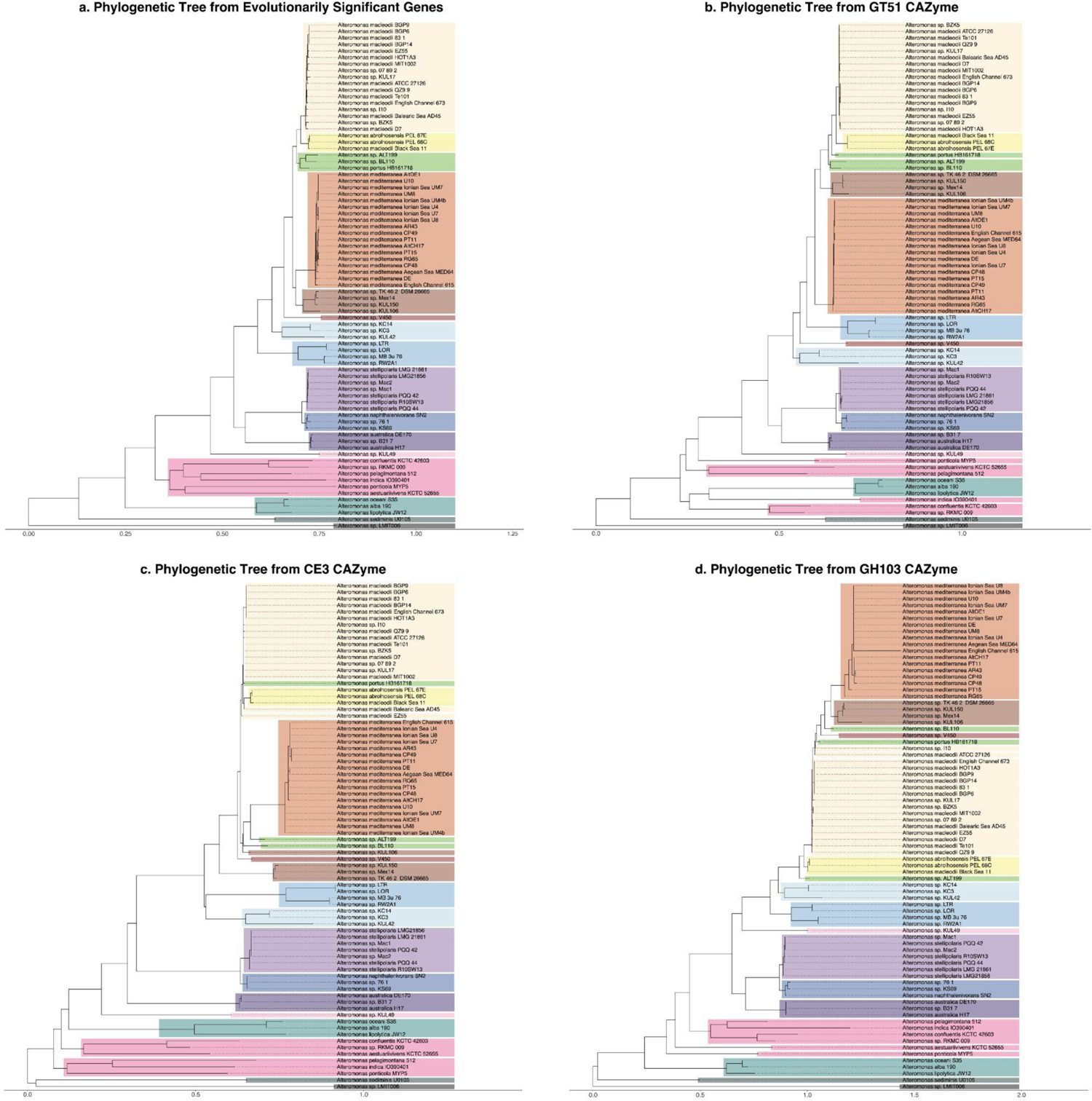
Individual phylogenies of core genes (a) and CAZymes (b, c, d) from the *Alteromonas* pangenome. The core genes (n=111) used to build the accepted phylogeny in (a) consist of those found in all 78 isolate genomes as single copy genes (‘--max-num-genes-from-each-genome’ set to 1), with the ‘--max-functional-homogeneity-index’ set to 0.9 and ‘--min-geometric-homogeneity-index’ set to 0.925. The CAZyme phylogenies (b, c, d) are each built from the alignment of one gene - GT51, CE3, and GH103, respectively. Clades are highlighted based on color assignments in Supplementary Figure 2. The horizontal axis provides a means for interpreting branch length as a measure of genetic change.

**Supplementary Figure 2.**
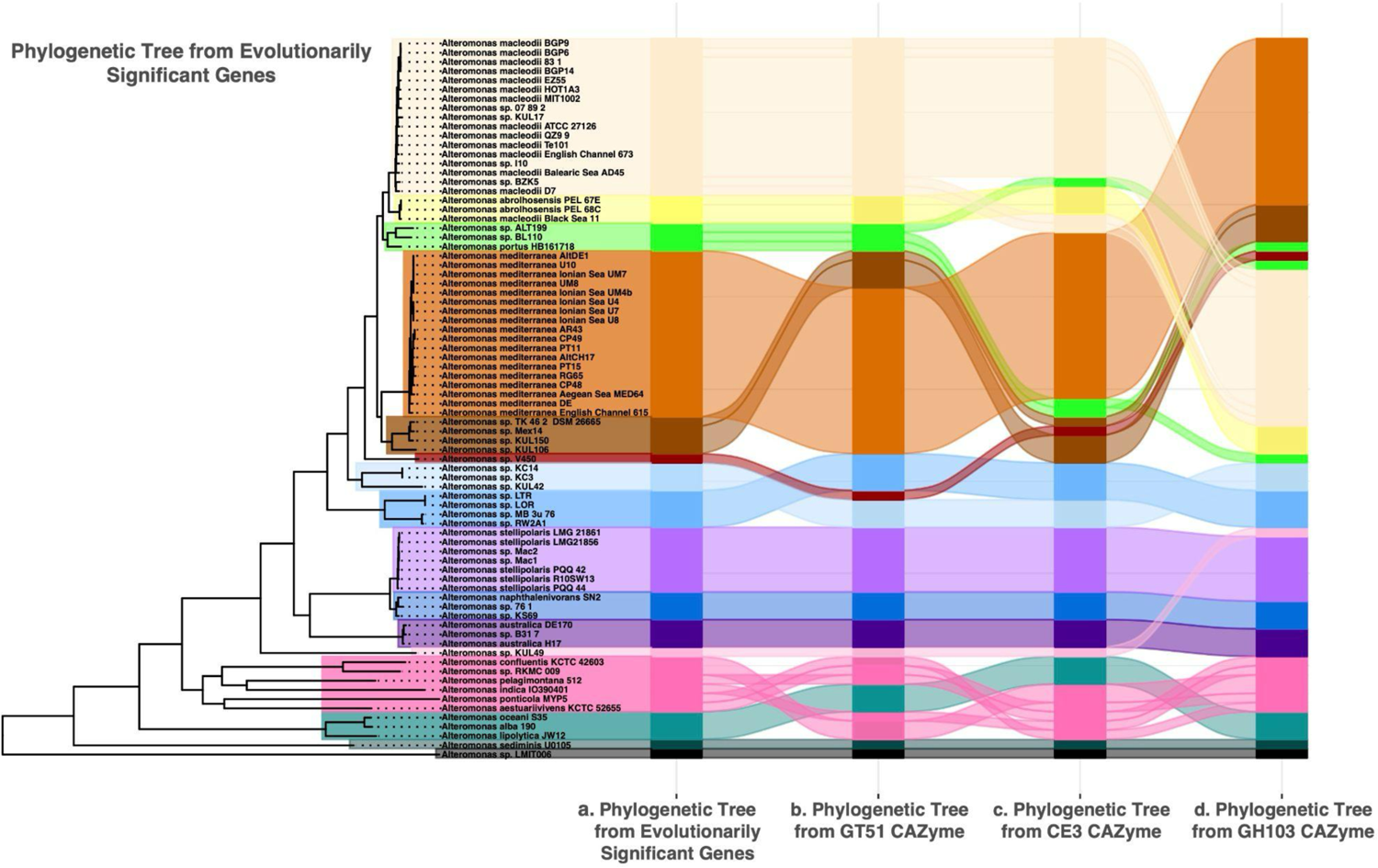
Comparison of core gene (a) and CAZyme (b, c, d) phylogenies for the *Alteromonas* pangenome. The core genes (n=111) used to build the accepted phylogeny in (a) consist of those found in all 78 isolate genomes as single copy genes (‘--max-num-genes-from-each-genome’ set to 1), with the ‘--max-functional-homogeneity-index’ set to 0.9 and ‘--min-geometric-homogeneity-index’ set to 0.925. The CAZyme phylogenies (b, c, d) are built from the alignment of one gene each. The alluvial plot displays the movement of clades within the tree. Further branch definition of each tree is shown in Supplementary Figure 1.

## Author Contributions

ZSC, MSS, LTR, SD, AME, MAM, and RB conceptualized the study. ZSC, MAD, IV, SM, CBS, LTR, WFS, MRM, PZL, and MS curated data. ZSC, MAD, MAM, and RB performed formal analyses. ZSC, MAD, CBS, LTR, WFS, MRM, PZL, and MS conducted investigations. IV, MSS, SM, and AME developed methodology. LW and AME administered the project. AME and MAM provided resources. IV, MSS, SM, and AME developed software. LW, AME, MAM, and RB supervised the project. ZSC, MAD, and IV validated the results. ZSC, MAD, LW, MAM, and RB worked on visualization. ZSC, MAD, IV, MSS, SM, LW, SD, AME, MAM, and RB wrote the paper with critical input from all authors.

